# Ultrafast and Ultralarge Distance-Based Phylogenetics Using DIPPER

**DOI:** 10.1101/2025.08.12.669583

**Authors:** Sumit Walia, Zexing Chen, Yu-Hsiang Tseng, Yatish Turakhia

## Abstract

**Motivation:** Distance-based methods are commonly used to reconstruct phylogenies for a variety of applications, owing to their excellent speed, scalability, and theoretical guarantees. However, classical *de novo* algorithms are hindered by cubic time and quadratic memory complexity, which makes them impractical for emerging datasets containing millions of sequences. Recent placement-based alternatives provide better algorithmic scalability, but they also face practical scaling challenges due to their high cost to compute evolutionary distances and significant memory usage. Current tools also do not fully utilize the parallel processing capabilities of modern CPU and GPU architectures.

**Results:** We present **DIPPER**, a novel distance-based phylogenetic tool for ultrafast and ultralarge phylogenetic reconstruction on GPUs, designed to maintain high accuracy and a small memory footprint. DIPPER introduces several novel innovations, including a divide-and-conquer strategy, a placement strategy, and an on-the-fly distance calculator that greatly improve the runtime and memory complexity. These allow DIPPER to achieve runtime and space complexity of *O(N.*log(*N))* and *O(N)*, respectively, with *N* taxa. With divide-and-conquer, DIPPER is also able to maintain a low memory footprint on the GPU, independent of the number of taxa. DIPPER consistently outperforms existing methods in speed, accuracy, and memory efficiency, and scales to tree sizes 1–2 orders of magnitude beyond the limits of existing tools. With the help of a single NVIDIA RTX A6000 GPU, DIPPER is able to reconstruct a phylogeny from 10 million unaligned sequences in under 7 hours, making it the only distance-based method to operate at this scale and efficiency.

**Availability:** DIPPER’s code is freely available under the MIT license at https://github.com/TurakhiaLab/DIPPER, and the documentation for DIPPER is available at https://turakhia.ucsd.edu/DIPPER. The test datasets and experimental results are available at https://zenodo.org/records/16803048.

## Introduction

Reconstructing phylogenies from molecular sequences is a fundamental problem in computational biology, underpinning a wide range of applications from evolutionary analysis to comparative genomics^1–3^. Among computational methods for this task, distance-based methods such as the Unweighted Pair Group Method with Arithmetic Mean (UPGMA)^4^, Neighbor-Joining (NJ)^5^, and Minimum Evolution (ME)^6^, although not the most accurate, offer a balance of speed and accuracy for many tasks. Some of these methods also provide strong theoretical guarantees^7,8^. Consequently, various distance methods-based tools have been developed and are widely used across a broad range of bioinformatics tasks, such as estimating a guide tree for multiple sequence alignments (MSAs)^9–16^, for bootstrapping or initializing heuristic trees for advanced phylogenetic methods involving Maximum likelihood (ML) and Bayesian inference^17–22^ for taxonomic classification of genomes^23,24^, and phylogenetic compression of molecular data^25^. Distance-based phylogenetic methods are also among the most widely used approaches for reconstructing phylogenies when MSAs are unavailable or impractical.

Classical *de novo* distance-based methods, particularly NJ and UPGMA, construct phylogenies by iteratively agglomerating taxa based on a pairwise distance matrix and have served as foundational methods in distance-based phylogenetics for decades. However, with the rapid surge in genome sequencing data, their scalability is becoming a significant bottleneck for some applications due to three key challenges associated with these algorithms. *First*, the agglomerative step incurs cubic time complexity (*O(N*^3^), where *N* is the number of taxa, rendering it impractical for large datasets. Tools such as RapidNJ^26^, CCPhylo^27^, FastPhylo^28^, QuickTree^29^, and DecentTree^30^ have been developed to mitigate this with implementation optimizations and *heuristics*, but have only extended the scalability to a few hundred thousand taxa. *Second*, these algorithms require a complete distance matrix as input, which imposes a quadratic space complexity (*O(N*^2^)), leading to impractical memory requirements for large datasets, which is often the dominant limiting factor, despite algorithmic speedups. *Third*, the computation of the distance matrix itself, especially for unaligned sequences, poses further challenges. While alignment-free methods for distance computation, such as MASH^31^, offer some relief, their runtimes for computing distance matrices are also impractical beyond a few hundred thousand sequences.

As *de novo* phylogenetic methods struggle to handle ultralarge datasets, phylogenetic placement approaches^6,32–44^, which insert a new taxon onto a fixed backbone tree, have emerged as a scalable alternative with a typical time complexity of *O(N*^2^). Examples of distance-based phylogenetic placement tools include FastME-2^36^ and APPLES^38^, which are based on balanced minimum evolution (BME) and least-squares (LS) criteria, respectively. APPLES-2^40^ introduced algorithmic enhancements based on a divide-and-conquer strategy, further reducing the time complexity to *O(N.*log*(N))*. Despite their scalability, these tools are bottlenecked by the cost of computing pairwise distances for unaligned sequences and the memory required to store distance matrices. For instance, while APPLES-2 partially alleviates the memory requirements by operating on an *M* x *N* matrix, where *M* is the number of queries and *N* is the number of backbone taxa, this approach is still impractical at extreme scales (with millions of sequences). Sparse Neighbor-Joining (SNJ)^43^ addresses the latter via recursive centroid-based partitioning and quartet resolution, reducing per-query memory complexity to *O(*log*(N))*. However, in the authors’ evaluations, this approach compromised accuracy and was not tested beyond 16K taxa, likely due to implementation inefficiencies.

To address the aforementioned limitations of existing distance-based phylogenetic methods, we introduce **DIPPER** (**DI**stance-based **P**hylogenetic **P**lac**ER**), a tool for reconstructing ultralarge phylogenies comprising up to tens of millions of taxa. DIPPER employs a novel *placement* algorithm that optimizes for the ME criterion, and employs a *divide-and-conquer* strategy to achieve runtime and memory complexities of *O(N.*log(*N))* and *O(N)*, respectively. DIPPER supports precomputed distance matrices or alignments as input, but also incorporates a GPU-accelerated ‘on-the-fly’ distance calculator using input MSA or raw sequences, allowing it to compute only the necessary pairwise distances during placement while maintaining a drastically low memory footprint. DIPPER’s placement strategy outperforms classical distance-based *de novo* approaches (e.g., NJ) on large datasets (10K-200K taxa), providing 44.4-731-fold higher speedup and up to 6.7-fold lower memory footprint at comparable or better accuracies. DIPPER also scales to ultralarge datasets, at least an order of magnitude beyond the capabilities of existing tools. It can reconstruct a tree from 1 million unaligned DNA sequences in 2 hours 21 minutes and from 10 million in just 6 hours 28 minutes—performance and scale not previously possible with distance-based phylogenetic methods.

## Methods

### DIPPER Overview

DIPPER is a scalable tool for distance-based phylogenetic tree reconstruction from unaligned or aligned nucleotide (DNA or RNA) sequences provided in FASTA format or using pairwise distance matrices in PHYLIP format^45^ (Fig. 1ai). In the inferred phylogeny, each tip is called a taxon (plural: taxa). Each taxon represents a single sequence from the input dataset. Therefore, the total number of taxa in the tree, *N*, is the same as the total number of input sequences. Depending on the input dataset size, DIPPER dynamically adapts its strategy between exact Neighbor-Joining, placement, and divide-and-conquer to optimize both performance and scalability (Fig. 1aii, methods), but users can also force a particular strategy by using a command-line parameter *--algorithm (-m).* For unaligned sequences, DIPPER computes pairwise distances on-the-fly using a GPU-accelerated implementation of the MASH algorithm^31^. This algorithm first converts each sequence into a *sketch* (a fixed-size vector consisting of the smallest hashed *k-mers*), then distances are computed via the Jaccard similarity metric computed with the help of the MinHash technique^46^ (see “Algorithmic Details”). This step enables fast, memory-efficient distance computation without the need for alignment.

For smaller datasets (fewer than 30K sequences), DIPPER employs the exact Neighbor-Joining (NJ) algorithm^5^, accelerated on GPUs for speed (Fig. 1aii, Methods). However, as datasets grow larger, the NJ algorithm becomes less practical due to its *O(N*^3^) time and *O(N*^2^) space complexity. To handle this, DIPPER adopts a novel placement algorithm for large datasets (30K–1M sequences), which incrementally builds the phylogeny under the minimum evolution (ME) criterion (Fig. 1b) and computes distances on-the-fly, reducing the time and space complexity to *O(N*^2^) and *O(N)*, respectively. The algorithm begins by constructing a backbone tree from two randomly selected sequences and then iteratively placing a query onto the growing tree. For each query, DIPPER computes (or retrieves) its distance to all current tips in the backbone tree. Each query is then evaluated for placement on all branches of the tree, with the optimal branch minimizing the increase in total tree length according to the ME criterion (Fig. 1b, see *Placement algorithm* in “Algorithmic Details”). To reduce computational overhead, DIPPER can restrict this evaluation to the *K* closest tips (default *K*=10) on each branch (e.g., *K*=4 in Fig. 1b), which are dynamically maintained throughout the process (see *K-closest placement algorithm* in “Implementation Details”).

**Figure 1:**
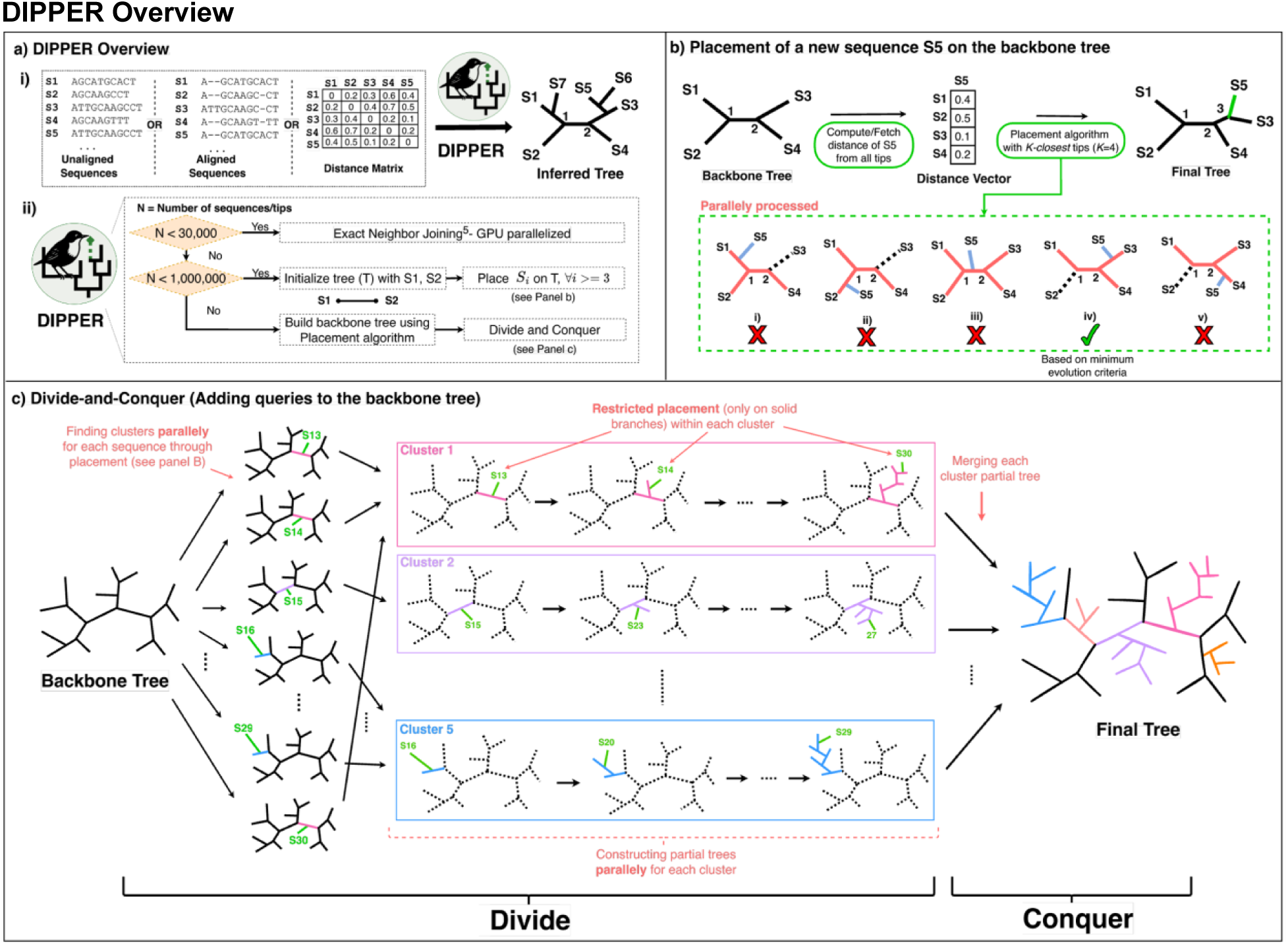
An overview of DIPPER. **(a) i)** DIPPER input and output, **ii)** DIPPER algorithm choice based on the number of taxa (N). If 𝑁 < 30𝐾, DIPPER reconstructs the tree using GPU-acceleration of exact Neighbor-Joining^5^, if 30𝐾 <= 𝑁 < 1𝑀, DIPPER builds the tree using its placement strategy based on minimum evolution criteria, and if 𝑁 >= 1𝑀, DIPPER uses a divide-and-conquer strategy on top of its placement strategy (Methods). **(b)** A detailed illustration of the placement strategy implemented in DIPPER. This algorithm adds a query sequence (S5 here) to a backbone tree by evaluating each branch in parallel and selecting the optimal branch based on the minimum evolution (ME) criterion. To minimize computational demands, DIPPER limits the evaluation up to the *K-closest* tips (*K*=4 here and default *K*=10), highlighted by colored branches. **(c)** A detailed illustration of the divide-and-conquer strategy in DIPPER. In the divide step, each new sequence is assigned to a distinct cluster based on the backbone branch it best aligns with under the minimum criterion. Subtrees are then independently built for each cluster (with possible deeper hierarchical levels not shown). In the conquer step, these subtrees are merged to form the final tree.

While this placement-based algorithm scales well for large datasets, ultralarge datasets (millions of sequences) still pose a challenge due to the algorithm’s *O(N*^2^) runtime. To handle this, DIPPER uses a divide-and-conquer strategy (Fig. 1c) that enables it to scale to an order of magnitude beyond existing tools. The process begins by constructing a backbone tree of a user-specified size *B* from a representative subset of sequences using the *placement algorithm*. The remaining sequences are then *clustered* based on the branch to which they are likely to attach instead of immediately placing them into the backbone. Then, sequences in each cluster are placed incrementally using a restricted placement scheme to construct partial trees. If a cluster exceeds the size *B*, it can be recursively subdivided. These partial trees are later merged to produce the final phylogeny. This divide-and-conquer algorithm achieves practical runtime and space complexity of *O(N.*log(*N))* and *O(N)*, respectively, significantly extending the scalability of phylogenetic inference (see Appendix A.1 in “Supplementary Materials”).

These algorithmic innovations are described in greater detail in the “Algorithmic Details” section. Beyond algorithms, DIPPER also incorporates novel GPU-aware optimization techniques (see “Implementation Details”) to fully leverage the massive parallelism within the tight memory budgets. Notably, its GPU memory usage is independent of the number of taxa (see Appendix A.1 in “Supplementary Materials”). Together, DIPPER’s algorithmic scalability, hardware-aware optimizations, and efficient implementation enable accurate reconstruction of phylogenies at unprecedented scales, surpassing the limits of current state-of-the-art tools in both speed and memory efficiency.

### Algorithmic Details

We now provide a detailed description of various algorithms adopted in DIPPER, which are illustrated in Fig. 1.

***Sketch Construction*:** This step is only required if the input is unaligned sequences. DIPPER employs the MinHash algorithm^46^ to convert unaligned sequences into fixed-size *sketches* of size *s*, i.e., *s-smallest* hashed *k-mers,* and store them in CPU (host) memory. It constructs a sketch for each sequence by converting all *k-mers* from the sequence into hashes and retaining only the *s* smallest hash values. Users can set the values of *s* and *k* using the command-line parameters *--sketch-size (-s)* and --*kmer-size (-k)*, respectively. By default, DIPPER uses *k=15* and *s=1000*.

***Exact Neighbor-Joining:*** Next, for small datasets containing fewer than 30,000 sequences, DIPPER employs the exact Neighbor-Joining algorithm to reconstruct the phylogenetic tree. If the distance matrix is not provided, DIPPER computes it from the input sequences (unaligned or aligned) using the distance calculator (see *Distance calculator* in “Algorithmic Details”).

***Placement algorithm:*** For larger datasets, containing up to a million sequences, DIPPER employs a novel placement algorithm to reconstruct the phylogeny. The goal of the placement algorithm is to determine the branch to insert a new taxon on a backbone tree while avoiding the underestimation of evolutionary distances in the resulting tree. This objective assumes that for non-additive matrices, the pairwise distance estimates of sequences represent lower bounds on the true evolutionary divergence between corresponding taxa. This objective can be replaced with the least squares criterion—widely used in prior studies^47,38,40^, without impacting the overall time complexity, but this criterion is more likely to violate the aforementioned assumption (see “Results”). Formally, DIPPER’s placement objective can be stated as:

Given a backbone phylogenetic tree *T* over a set of taxa *X*, a query taxon *t*, and a distance vector δ(i, t) for each *i ∈ X,* insert *t* on a branch *(u*, v*)* of *T* such that the resulting tree *T′* satisfies *d(i, t) ≥ δ(i, t)* for all *i* in the set 𝐿_𝑘/2_ ^𝑢∗→𝑣∗^ and 𝐿_𝑘/2_ ^𝑣∗→𝑢∗^, where *d(w, t)* denotes the evolutionary distance between a node *w* and the query taxon *t* in the resulting tree (see Supplementary Fig. 1 for an illustrative explanation), and 𝐿_𝑘_^𝑝→𝑞^ denotes the *k*-*closest* tips to node *p* whose shortest paths to *p* do not traverse through *q,* while resulting in a minimum additional branch length 𝛥(𝑢 ∗, 𝑣 ∗; 𝑡) (as per the ME criterion).

Mathematically,

𝑑(𝑖, 𝑡) >= 𝛿(𝑖, 𝑡) ∀ 𝑖 ∈ 𝐿_𝑘/2_^𝑢→𝑣^ Avoid underestimations

Since 𝑑(𝑖, 𝑡) = 𝑑(𝑢, 𝑡) + 𝑑(𝑢, 𝑖) ∀ 𝑖 ∈ 𝐿_𝑘/2_^𝑢→𝑣^ (see Supplementary Fig. 1)

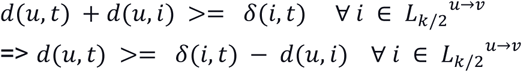

Hence,

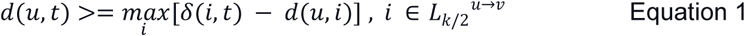

Similarly,

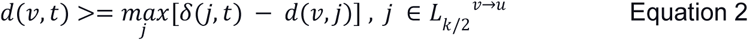

The additional branch length 𝛥(𝑢, 𝑣; 𝑡), incurred by placing *t* on the branch *(u,v),* is described by

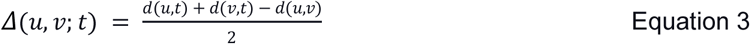

Since 𝑑(𝑢, 𝑣) is already fixed in the tree, therefore, 𝛥(𝑢, 𝑣; 𝑡) in equation 3 can be minimized when the LHS of equations 1 and 2 are simultaneously minimized, i.e., when:

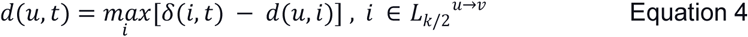

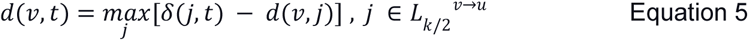

Therefore, the best placement branch (𝑢 ∗, 𝑣 ∗) according to the ME criterion is the one among all possible *(u,v)* ∈ *T* that minimizes the additional branch length.

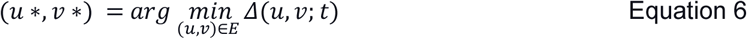

DIPPER does not allow negative branch lengths, so the value of 𝛥(𝑢 ∗, 𝑣 ∗; 𝑡) is set 0 if found negative. DIPPER’s *K-closest* algorithm (described above) for placement of one sequence on a tree of *n* taxa has a time complexity of *O(Kn),* as each placement evaluates *O(n)* branches in the tree (equation 6), each using distance estimates from its *O(K)* nearest nodes (equations 4 and 5). Therefore, when building a tree of *N* taxa from scratch, the algorithm has a time complexity of *O(KN*^2^), which simplifies to *O(N*^2^) since *K* is a user-defined constant. DIPPER’s placement strategy significantly improves scalability and mitigates the underestimation of evolutionary distances prevalent in earlier approaches (see “Results”). Users may disable the *K-closest* heuristic by setting *K=-1* through the command-line argument *--K-closest (-K)*, enabling exhaustive evaluation across all tips and eliminating evolutionary distance underestimation (see “Results”).

By avoiding the need to recompute global topologies, computing distances (δ(i, t) ∀ 𝑖 ∈ 𝑋) on-the-fly, DIPPER’s placement algorithm offers significant advantages in both speed and memory efficiency, while maintaining the accuracy, enabling it to scale up to a million sequences.

***Divide-and-Conquer Strategy:*** For ultralarge datasets, containing more than a million sequences, DIPPER adopts a divide-and-conquer strategy. We now describe the four steps involved in this algorithm (Fig. 1c). Step (a) is an *initialization* stage, steps (b) and (c) together constitute the *divide* stage, and step (d) is the final *conquer* stage.

*a) Backbone Tree Construction:* To initiate the divide-and-conquer strategy, DIPPER first constructs a backbone tree from a random subset of sequences. In DIPPER, the default value for the backbone tree is set to 100K, and can be set by the user through a command-line argument *--backbone-size (-B)*. This backbone is built using the *K-closest placement algorithm* (see “Implementation Details”), where each taxon is placed onto a growing tree based on its computed distances to existing taxa under the minimum evolution criterion. The goal is to capture the broad structure of the tree while keeping the initial construction computationally feasible. The resulting backbone serves as a scaffold for the subsequent placement of the remaining taxa. The runtime complexity of this step for a backbone of *B* taxa is *O(B*^2^). Note that *B* is a user-specified constant, independent of *n*, the total number of taxa provided.
*b) Clustering Based on Predicted Placement:* Once the backbone tree is built, the remaining taxa, those not included in the initial backbone, are assigned to clusters based on their predicted branch of attachment to the backbone. For each query sequence, DIPPER computes or fetches a distance vector to the tips of the backbone and determines its most likely placement branch. Rather than immediately incorporating these taxa into the backbone, they are grouped into clusters corresponding to the branch of the backbone where they are likely to belong based on the *K-closest placement algorithm*. This clustering enables localized and parallelizable tree construction in the next step. By default, DIPPER sets the maximum cluster size equal to the size of the backbone tree (i.e., same as *B*). Users may override the default by specifying a custom maximum cluster size using the command-line argument *--max-cluster-size (-C)*. If a cluster exceeds this limit, the algorithm recursively constructs the backbone trees and subdivides the cluster until all resulting clusters meet the size constraint.
*c) Partial Tree Construction:* Within each cluster, DIPPER incrementally constructs a partial tree (subtree). Sequences in a cluster are placed and incorporated using a *restricted placement scheme*—a constrained version of the *K-closest placement* approach that limits insertion within a predicted backbone branch. This restriction reduces computational overhead. Each cluster is processed independently, allowing for parallel construction of partial trees on GPUs. Since the maximum size of the cluster is determined by a user-specified constant, each partial tree can be constructed on a GPU in constant time and memory, independent of the total number of input taxa. This is an important property that allows DIPPER to accelerate the computation of ultralarge trees even on memory-constrained GPU devices.
*d) Merging Partial Trees:* Finally, after all clusters have been processed and their partial trees constructed, DIPPER performs a merging step to integrate these subtrees back into the global phylogeny. Each partial tree is grafted onto the original backbone at its corresponding attachment point, preserving both the high-level structure of the backbone and the finer resolution (including the branch lengths) provided by the local subtrees. This final merge yields the complete phylogenetic tree that incorporates all input sequences at its tips (Fig. 1c).

This divide-and-conquer strategy has a total expected runtime complexity of *O(N.*log(*N))* (see Appendix A.1 in “Supplementary Materials” for proof).

***Distance calculator*:** DIPPER’s distance calculator computes a distance vector, representing the distances between a query sequence and a set of target taxa, which form the tips of a backbone tree, from either *sketches* or input alignments. When using *sketches*, distances are computed via the Jaccard similarity metric, following the approach used in MASH^31^. For alignments, DIPPER supports evolutionary distance estimation using models such as Jukes–Cantor (JC69)^48^, Tajima–Nei (TN93)^49^, Kimura two-parameter (K2P)^50^, and Tamura^51^. Users can choose the evolution model in DIPPER software using the command-line parameter *--distance-type (-t)* to estimate the distances from alignments; the JC69 model is used by default. If the distance matrix is provided, the kernel fetches the necessary vector. This method is particularly useful when placing a query onto a backbone tree, as it reduces the space complexity to *linear* (with the number of taxa) compared to *quadratic* in most previous works.

### Implementation Details

DIPPER is implemented in C++ and leverages Threading Building Blocks (TBB)^52^ to fully exploit thread-level parallelism available on multi-core platforms, with key computational kernels to be accelerated on GPUs implemented using CUDA. Our implementation incorporates several optimizations to support efficient parallel processing and minimize memory overhead. These include fine-grained thread-block level parallelism, dynamic memory management strategies, batching, and the use of shared memory primitives to reduce latency and maximize throughput. Together, these design choices enable DIPPER to efficiently scale to millions of sequences on modern GPU architectures.

***K-closest placement algorithm:*** To reduce the computational cost associated with evaluating candidate placements, DIPPER implements a *K-closest placement* strategy that heuristically limits the number of comparisons required per branch. For a given query sequence and a candidate branch *(u,v)*, instead of comparing against all descendant tips of *u* and *v*, DIPPER considers only the *K/2* nearest tips from *u* and *v* (Equations 1 and 2). These closest tips are used to approximate the contribution of each tip to the path length difference incurred by placement (𝛥 in Equation 3). The set of *K-closest* tips is dynamically updated as new taxa are added to the backbone, ensuring that the approximation remains robust throughout the tree construction process. Empirically, this heuristic maintains placement accuracy—often matching or exceeding that of exhaustive evaluation—while offering substantial improvements in runtime performance, especially on large phylogenies (see Results). In DIPPER, the default value for *K* is set to 10, and can be set to *-1* through the command-line argument *--K-closest (-K)* to perform strict mode, i.e., considering all taxa for the comparison.

***GPU Parallelism techniques:*** DIPPER leverages both coarse-grained and fine-grained parallelism for GPU acceleration across all its algorithmic stages to enable rapid and scalable phylogenetic construction. The following strategies are employed across its GPU kernels:

a. **Sketch construction kernel:** DIPPER accelerates the sketch construction step using a two-level GPU parallelism strategy. At the coarse level, each input sequence is processed independently by assigning one thread block per sequence. Within each block, fine-grained parallelism is used to process the first *s* consecutive *k-mers*, with each thread hashing one *k-mer* and contributing to a parallel sort to initialize the sketch. To perform sorting, DIPPER uses the radix sort implementation from the NVIDIA CUDA CUB library (https://docs.nvidia.com/cuda/cub/index.html), which operates directly in shared memory. This allows extremely fast, in-place sorting of hash values with no need for global memory access, contributing to the overall speed and efficiency of sketch construction. In subsequent iterations, each block processes the next *s k*-mers, hashes them, and merges them with the existing sketch using another sort step that retains only the *s* smallest hash values. This iterative update continues until the entire sequence is processed, yielding a fixed-length sketch per sequence. A key optimization in DIPPER is the use of shared memory to maintain only 2*s* hash values per block at any time. This significantly improves memory efficiency and minimizes access to slower global memory, which is especially beneficial when scaling to large numbers of sequences. Because shared memory is much faster and has lower latency, storing intermediate hash values in this space allows efficient in-block processing without the overhead of global memory traffic.
b. **Distance Computation kernel:** The pairwise distance computation kernel in DIPPER is designed to scale to ultralarge datasets with linear space complexity (*O(N)*) with the number of sequences. When required distance vectors to certain sketches or alignments are not already on the GPU, they are transferred from host memory, when available at the host, or computed on-the-fly on the GPU. The GPU-accelerated kernel capitalizes on the high throughput of the hardware and the independence of distance evaluations, supporting efficient and parallel computation across large collections of taxa. Each element of the distance vector is computed independently across GPU threads, with thread-block-level parallelism enabling simultaneous calculation of all pairwise distances.
c. **Placement kernel:** For each query sequence, DIPPER’s placement algorithm evaluates all candidate branches on the reference tree in parallel. Each candidate branch is assigned to a separate GPU thread, which computes the associated increase in branch length and stores it in a shared vector. The optimal placement is then identified using a parallel reduction operation (using the NVIDIA Thrust library, https://developer.nvidia.com/thrust), which efficiently determines the minimum added branch length. Query taxa are added sequentially, but within each placement step, the exhaustive evaluation of branches is massively parallelized. This design becomes increasingly beneficial as the tree size grows, allowing tens of thousands of branches to be evaluated concurrently.
d. **Divide-and-Conquer kernel:** DIPPER further exploits the high degree of parallelism in its divide-and-conquer pipeline. Key stages, including *backbone tree construction*, *clustering based on predicted placement*, and *partial tree construction,* are parallelized across GPU threads and thread blocks, exploiting the inherent parallelism associated with its core *K*-closest placement *algorithm*, which is central to DIPPER’s scalability and accuracy in processing large-scale phylogenomic datasets.

***Memory efficiency techniques:*** Distance-based phylogeny reconstruction is inherently memory-intensive, posing a major challenge to the scalability of most state-of-the-art tools, even on modern systems. DIPPER addresses this limitation through the following techniques.

a. **On-the-Fly Distance Calculation:** Conventional phylogenetic reconstruction tools compute and store the full pairwise distance matrix, incurring a quadratic space complexity, *O(N*^2^), with the number of input taxa. DIPPER circumvents this bottleneck by computing distances dynamically, on-the-fly, during phylogenetic placement. In both its placement and divide-and-conquer kernels, DIPPER places each query sequence onto a backbone tree. Importantly, this operation requires only the distances between the query and the tips of the backbone, not the full distance matrix (Fig. 1b). By computing this distance vector on demand, DIPPER reduces the memory requirement from quadratic to linear with respect to the number of backbone tips. To the best of our knowledge, no existing tool applies this strategy when reconstructing phylogenies from unaligned sequences, limiting their scalability on ultralarge inputs. This optimization is one of the key enablers of DIPPER’s ability to handle datasets at the scale of tens of millions of taxa.
b. **Multi-Level Batching for Memory-Constrained Processing:** DIPPER employs a two-level batching strategy to address memory constraints at both the CPU and GPU levels.

I. **Input Batching (CPU Memory Management):** When processing datasets with millions of DNA sequences, storing the complete input—especially aligned sequences—can exceed the capacity of the CPU’s main memory. DIPPER addresses this by processing the input in batches. Each batch contains only as many sequences as can fit in memory and is immediately compressed: unaligned sequences are encoded using a 2-bit and 4-bit representation for unaligned and aligned sequences, respectively. Ambiguous characters are converted to the DNA/RNA character ‘A’, similar to the MASH algorithm. These compressed batches are stored and reused as needed throughout subsequent stages. This design allows DIPPER to manage memory efficiently and avoid loading the entire dataset into memory at once.
II. **GPU-level Batching:** DIPPER computes the distance between each query and all tips of the backbone tree efficiently on the GPU. However, due to limited GPU global memory, it is often not feasible to store all backbone tip sequences—even in compressed form—on the device. A naive workaround involving repeated transfers of data batches between CPU and GPU for each query results in significant overhead, negating the speed advantages of GPU acceleration. DIPPER overcomes this problem through its divide-and-conquer strategy. In this approach, the backbone tree is designed to be small enough for its tip sequences to fit entirely into GPU memory. The backbone remains fixed during the clustering step, allowing DIPPER to batch multiple queries to the GPU efficiently and compute distances with minimal transfer overhead. In subsequent steps, where partial trees are constructed for constrained placements, the memory requirements are even lower. This hierarchical processing ensures that each GPU-bound operation is both memory-efficient and computationally effective.

Together, these techniques—dynamic distance computation, input-level compression, and GPU-aware batching—enable DIPPER to scale to previously intractable dataset sizes. Using these strategies, DIPPER has successfully reconstructed phylogenies for datasets containing up to 10 million taxa using 224GB and 23GB of CPU and GPU memory, respectively, demonstrating its practical scalability and efficiency.

### Experimental Methodology

**Dataset:** We evaluated DIPPER and baseline tools using three simulated datasets of diverse characteristics and scale. 1) The first dataset was simulated using AliSim^53^, based on the General Time Reversible (GTR) model, with an average branch length of 0.05 substitutions per site and an indel rate of 0.12 indels per substitution. This dataset includes trees with taxa counts ranging from 10,000 to 1,000,000 and sequences with an average length of 10,000 bases. 2) The second simulated dataset was generated using RNASim^54^, in which taxa were randomly sampled in sizes ranging from 10,000 to 1,000,000 from a reference tree containing one million sequences. The average sequence length in the RNASim dataset is approximately 1,500 bases. 3) Finally, we used AliSim to simulate the largest dataset containing 10 million sequences, using a random tree simulated under the Yule-Harding^55^ speciation model, based on the GTR model with an average branch length of 0.002 substitutions per site and an indel rate of 0.12 indels per substitution. For all three datasets, the trees used for simulation were treated as the true trees for accuracy evaluation.

**Baseline Tools:** We used the latest available versions of RapidNJ^26^, FastME^36^, QuickTree^29^, DecentTree^30^, CCPhylo^27^, FastPhylo^28^, and APPLES-2^40^ as baselines to compare with DIPPER. We also attempted to compare with SNJ^43^, but could not get it to run successfully. The remaining tools were executed with default parameters. For DecentTree, we opted for the vectorized Neighbor-Joining (NJ-V) algorithm because it is the fastest and most scalable option available. We used MASH^31^ to compute the distance matrix from unaligned sequences for all baseline tools. The only exception is FastPhylo, for which we utilized the fastdist^28^ program to estimate distances, as specified by the authors. All tools were executed under the Jukes-Cantor^48^ (JC69) model of evolution to estimate trees from alignments, except CCPhylo and QuickTree, for which we used the default mode, as these tools do not allow users to select a specific model for their analysis. We used TWILIGHT^16^ to construct multiple sequence alignments using guide trees provided by various tools in comparison. Finally, we also compared DIPPER to a recent K-medoids clustering algorithm introduced in FAMSA-2^56^ for guide tree construction of large MSAs. See Appendix A.2 in “Supplementary Materials” for detailed information on software versions and commands used for all baselines.

**Metrics:** To evaluate the accuracy of estimated trees, we used MAPLE 0.6.12^42^ to compute the normalized Robinson-Foulds distance (nRF) of the inferred tree to the true tree from the simulator for the simulated dataset. We used GNU time (https://www.gnu.org/software/time/) to measure the runtime of all tools. We measured peak memory usage using `VmPeak` from Linux */proc* files and NVIDIA Nsight Compute (https://docs.nvidia.com/nsight-compute/) for CPUs and GPUs, respectively.

**Hardware and Execution Environment:** DIPPER and all baseline tools were executed on NSF Expanse, an Advanced Cyberinfrastructure Coordination Ecosystem: Services and Support (ACCESS) cluster. Each Expanse CPU node has two 64-core AMD EPYC 7742 processors with 256 GB of DDR4 memory, and a GPU node has four NVIDIA V100s (32 GB SMX2) connected via NVLINK and dual 20-core Intel Xeon 6248 CPUs. We used one CPU node and one GPU node. All experiments, unless otherwise specified, were conducted using 32 CPU cores and 1 GPU on the parallelized versions of different tools.

## Results

### DIPPER provides excellent speed and accuracy for building ultralarge phylogenies

We evaluated DIPPER’s speed and accuracy with state-of-the-art distance-based phylogeny reconstruction tools by constructing trees from unaligned sequences generated using AliSim (Dataset 1, see “Experimental Methodology”). For a fair comparison, if a baseline tool did not include distance computation from unaligned sequences, we included the runtime of MASH to the tool’s total runtime, which was used to provide it with the required input distance matrix (Fig. 2). All tools were allowed to run for a maximum 24-hour time limit for each input dataset.

**Figure 2:**
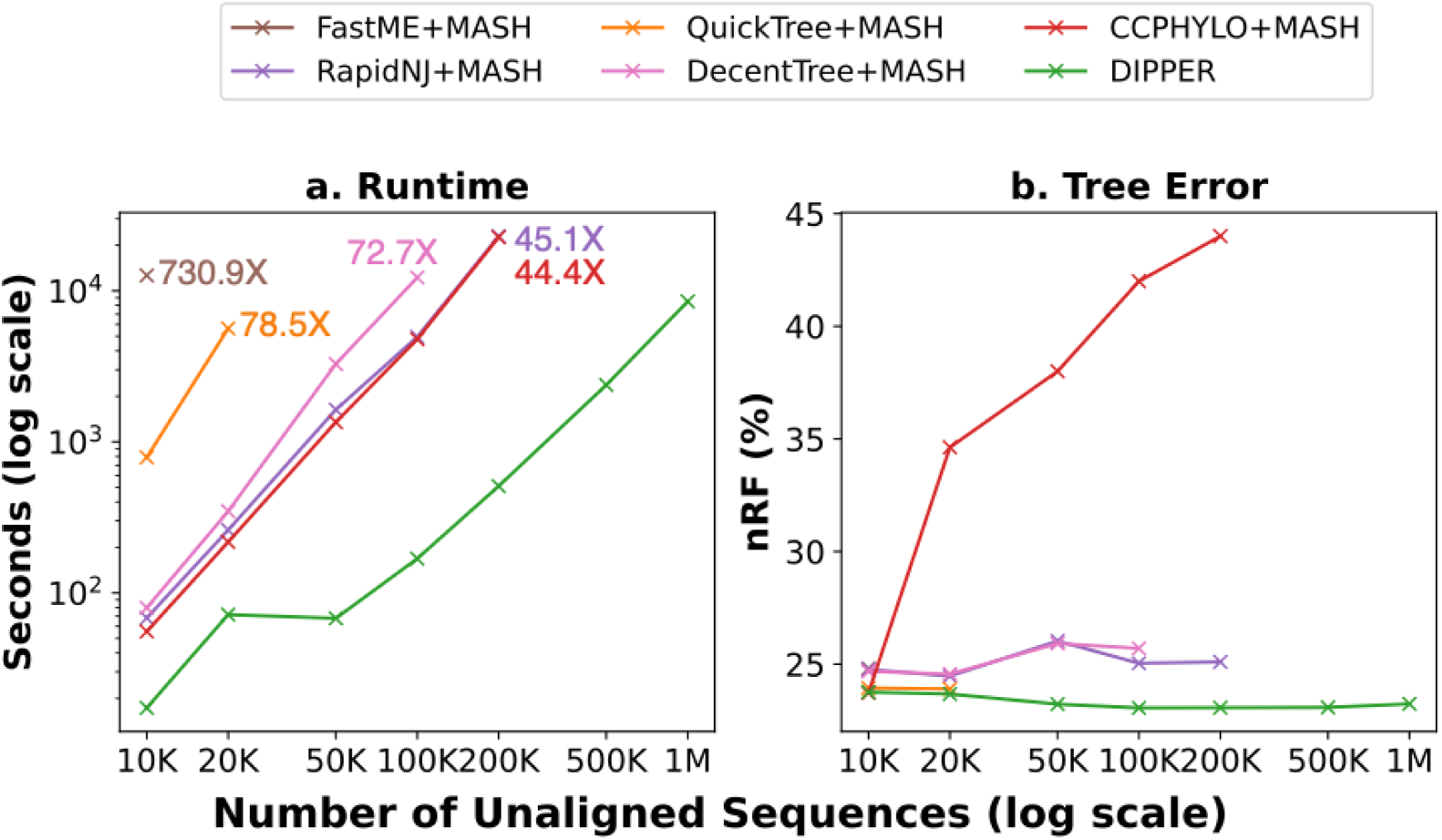
Runtime and accuracy with unaligned sequence count. **(a)** Runtime and **(b)** Accuracy of DIPPER compared with state-of-the-art tools to construct phylogenies from unaligned sequences generated using AliSim. We provide the ratio of the runtimes of different tools relative to DIPPER on the largest dataset they could handle.

DIPPER stands out to be the fastest and most accurate among all evaluated tools (Fig. 2). DIPPER is the only tool capable of constructing a tree from 1 million unaligned sequences. It provides 44.4-fold and 45.1-fold speedup against the closest competitors, CCPHYO and RapidNJ, at 200,000 sequences, and between 72.7-fold and 730.9-fold compared to the remaining tools. FastME and QuickTree failed to reconstruct the phylogeny for 20,000 and 50,000 sequences, respectively, within the time limit of 24 hours. Other baseline tools were limited by their memory utilization and the runtime of MASH, which scales up to a dataset with 100,000 sequences and fails to estimate the distance matrix for larger datasets within the time limit. DIPPER scales better than all state-of-the-art tools. As the dataset size increases to 50,000 sequences and beyond, DIPPER switches to a placement-based approach, as opposed to exact NJ for smaller datasets (10,000 and 20,000 sequences), resulting in a substantial speedup. Notably, it reconstructs a tree for 50,000 taxa in just 68 seconds, comparable to the time it takes to reconstruct a phylogeny for 20,000 taxa using NJ. Along with the GPU-accelerated placement strategy, the on-the-fly distance computation kernel helps DIPPER to reconstruct the phylogeny for 1 million sequences in just 2 hours and 21 minutes.

In terms of accuracy, DIPPER consistently matches or outperforms all evaluated tools. For the dataset with 10,000 sequences, all methods achieve similar nRF error rates (23.3%-24.8%). As the dataset size increases, CCPhylo’s accuracy deteriorates and continues to be the least accurate across all scales as it uses a heuristic NJ. The error rates for the remaining tools remain relatively stable, with DIPPER emerging as the most accurate. At 100,000 sequences, DIPPER achieves a 2% and 2.6% lower nRF error rate than RapidNJ and Decentree, respectively, indicating that DIPPER’s placement heuristic is more accurate than classical NJ on these datasets (Fig. 2). DIPPER’s accuracy improves with growing datasets, reaching 23.2% error at 1 million sequences—further reinforcing its robustness in large-scale distance-based phylogenetic reconstruction.

### DIPPER can perform ultralarge distance-based phylogenetic reconstruction from input alignments

To assess the performance of DIPPER, we benchmarked its speed and accuracy against state-of-the-art distance-based phylogenetic reconstruction tools, using aligned sequences generated by RNASim (Dataset 2, see “Experimental Methodology”). Across all tested scenarios, DIPPER emerged as both the fastest and most scalable tool, while maintaining comparable or better accuracy than the baseline tools (Fig. 3).

**Figure 3:**
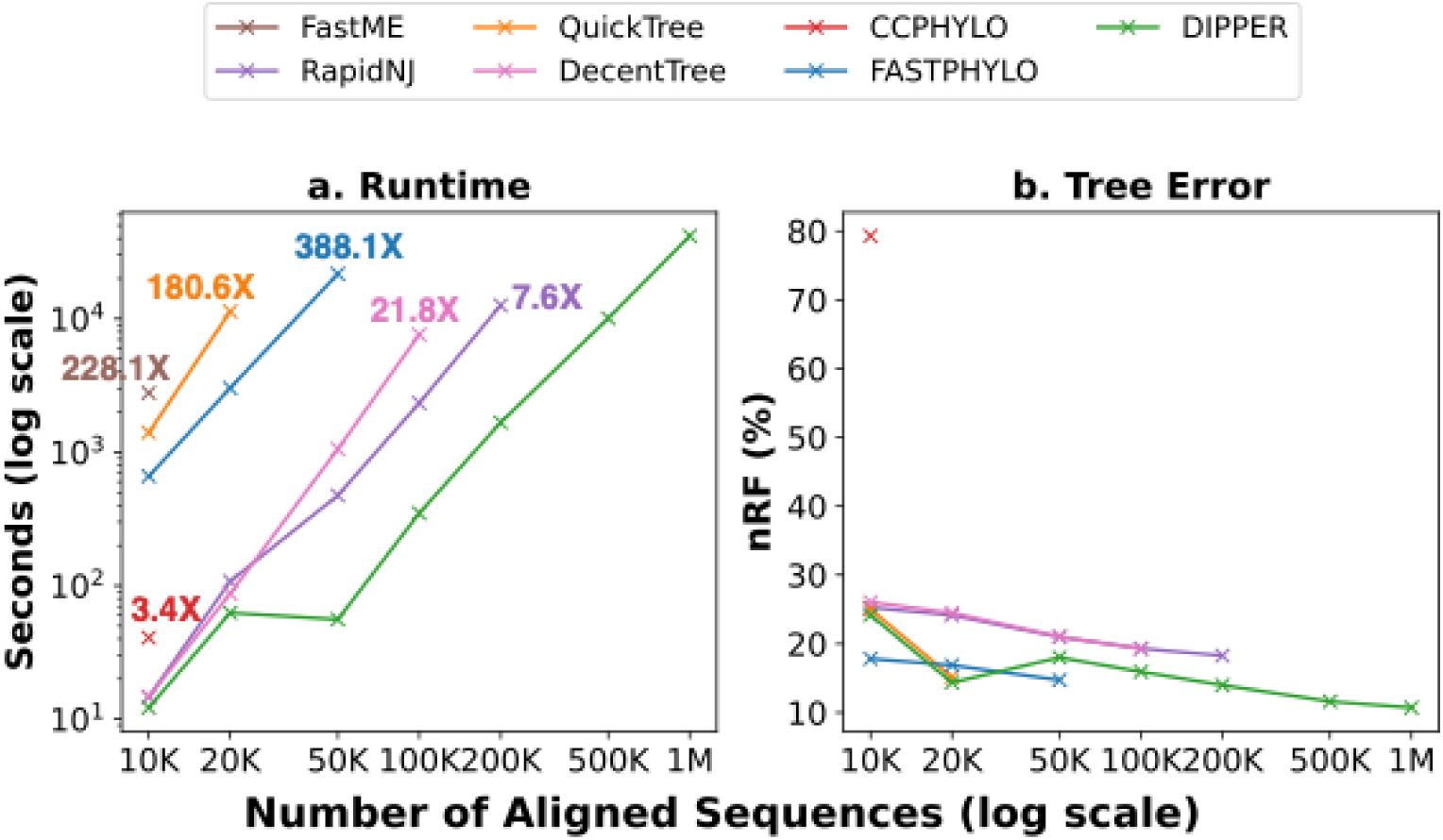
Runtime and accuracy with aligned sequence count. **(a)** Runtime and **(b)** Accuracy of DIPPER compared with state-of-the-art tools to construct phylogenies from aligned sequences generated using RNASim. We provide the ratio of the runtimes of different tools relative to DIPPER on the largest dataset they could handle.

Again, DIPPER is the only method among those evaluated capable of constructing a phylogenetic tree from one million aligned sequences. At 200,000 sequences, it achieves a 7.6-fold speedup over RapidNJ, its closest competitor, and demonstrates speed improvements ranging from 3.4-fold to 388.1-fold compared to the other tools. A few of the baseline tools demonstrate scalability issues with small datasets. For instance, FastME and QuickTree failed to complete the run within the 24-hour time limit beyond 20,000 and 50,000 sequences, respectively, and CCPhylo’s distance calculator terminated with a “*No sufficient overlap was found*” error for datasets larger than 10,000 sequences. Other tools were constrained by memory usage, with several exceeding 256GB of RAM available due to the need to load the complete distance matrix in main memory. DIPPER scales better than all state-of-the-art tools. With the *K-closest placement algorithm* and an efficient on-the-fly distance computation kernel, DIPPER reconstructs the phylogeny from one million sequences in just 11 hours and 45 minutes, using 92.3 GB of memory and staying within practical memory limits.

In terms of accuracy, DIPPER performs competitively across all tested scales. It consistently matches or exceeds the accuracy of all tools except FastPhylo, which implements a heuristic for NJ. However, FastPhylo does not scale beyond 50,000 sequences, limiting its practical utility. For the 10,000-sequence dataset, all tools (except CCPhylo) report comparable normalized Robinson-Foulds (nRF) error rates ranging between 24.1% and 26.1%. Notably, on smaller datasets, DIPPER performs exact NJ, thus matching the accuracy of QuickTree. DIPPER’s accuracy improves as the dataset grows: despite a slight increase in error at 50K taxa when shifting from exact NJ to the *K-closest* placement strategy, DIPPER also maintains better accuracy than DecentTree and RapidNJ at comparable scales. Crucially, as the dataset size increases, DIPPER’s error rate steadily decreases (which is also true of most other baselines, Fig. 3b). Specifically, it achieves a reduction in nRF error from 24.1% at 10,000 sequences to 10.7% at one million sequences. This trend underscores DIPPER’s effectiveness and reliability for large-scale distance-based phylogenetic reconstruction from aligned data.

### DIPPER builds ultralarge phylogenies with low memory requirements

DIPPER’s on-the-fly distance computation, bypassing the need to estimate and store a complete distance matrix, enables efficient reconstruction of ultralarge phylogenies with substantially reduced memory requirements. We assessed DIPPER’s memory consumption and benchmarked it against state-of-the-art tools through constructing phylogenies from unaligned sequences simulated with AliSim (Dataset 1, see “Experimental Methodology”). For DIPPER, we report the combined memory usage of both CPU and GPU resources to reflect the total system footprint.

For larger datasets, DIPPER exhibits the most memory-efficient profile among all evaluated tools. Theoretically, the memory usage of DIPPER should scale linearly (*O(N)*) with the number of sequences, but, empirically, its memory usage scales sublinearly with the number of input sequences, following an approximate *O(N*^0.4^) growth, in sharp contrast to the *O(N²)* memory complexity typical of traditional distance-matrix-based approaches (Fig. 4). For smaller datasets (10,000–20,000 sequences), memory consumption across tools is generally low, with DIPPER using more memory than some baselines. However, memory usage quickly becomes a limiting factor for other methods: QuickTree and FastME fail to process 20,000 and 50,000 sequences, respectively, while DecentTree exceeds our available memory (256 GB) beyond 100,000 sequences, requiring 6.7-fold more memory than DIPPER at that scale. CCPhylo and RapidNJ manage to scale up to 200,000 sequences but consume 146 GB of memory, over 5.7-fold higher than DIPPER, making them impractical for much larger datasets. In contrast, DIPPER maintains a sub-linear increase in memory usage, growing from 11.6 GB at 10,000 sequences to just 86.1 GB at 1 million sequences, comfortably within the limits of modern high-performance computing infrastructure. For all datasets, the GPU memory usage of DIPPER was within 10GB, also comfortably within the limits of modern GPUs.

**Figure 4:**
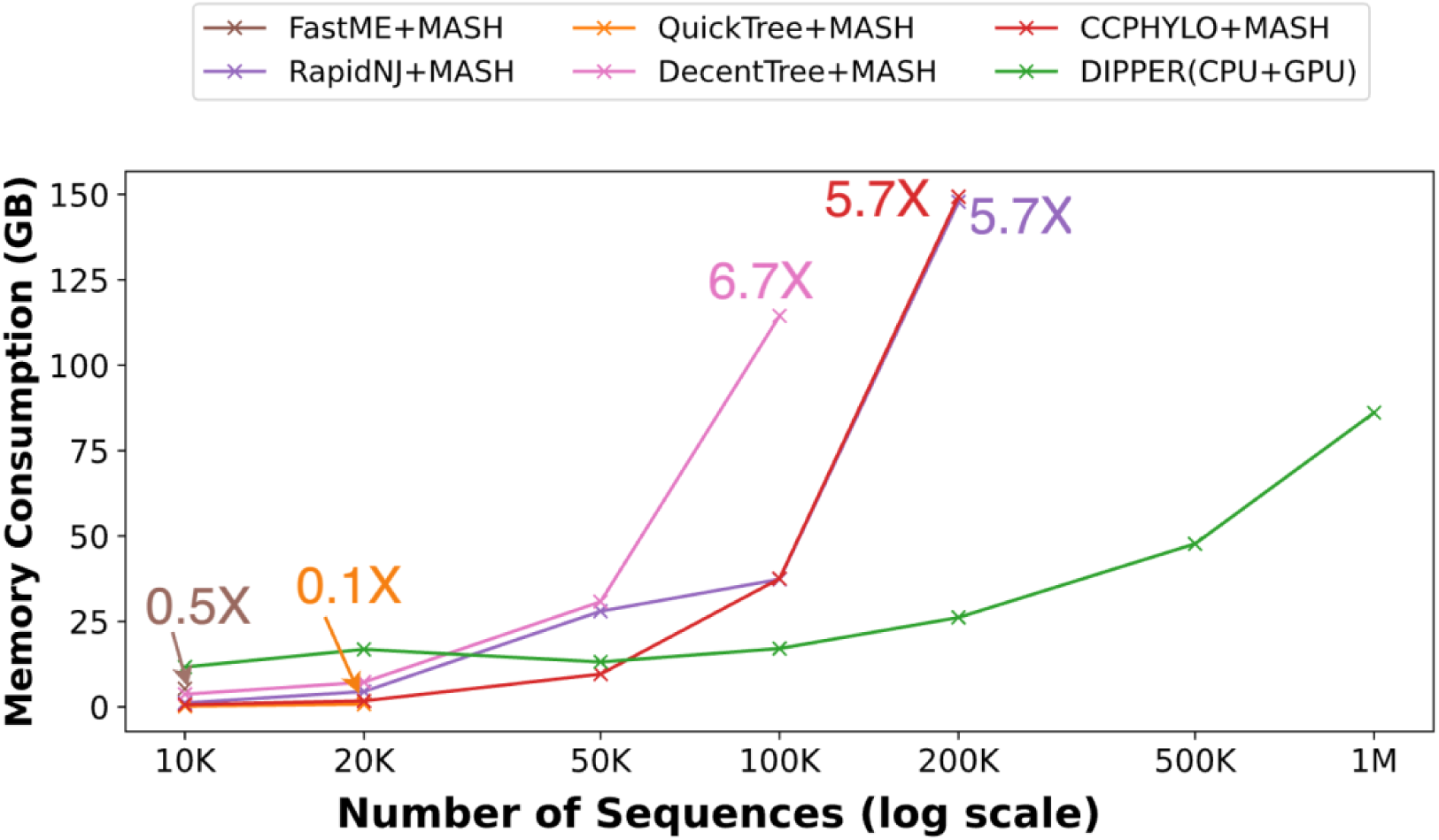
Memory usage with sequence count. Memory usage of DIPPER compared with state-of-the-art tools. We report the combined memory usage of both CPU and GPU resources to reflect the total system footprint of DIPPER.

### DIPPER accelerates the construction of ultralarge alignments

Progressive alignment is one of the most widely used strategies for constructing MSAs. In this approach, sequences are aligned sequentially based on their positions in a *guide tree*, starting with the most closely related pairs and moving toward more distantly related ones. This method underlies many scalable MSA tools^9–11,14–16^; however, its effectiveness is fundamentally constrained by the quality and scalability of the guide tree. TWILIGHT^16^ is a state-of-the-art progressive aligner that has demonstrated the potential to construct ultralarge MSAs, containing millions of sequences, with both high accuracy and speed. Yet TWILIGHT’s scalability is limited by its dependence on external guide trees, for which no method exists to operate efficiently at such a scale.

DIPPER directly addresses this limitation by enabling guide tree construction for ultralarge datasets. In combination with TWILIGHT, DIPPER enables the alignment of one million unaligned sequences in just 3 hours and 2 minutes, representing a major advance in both performance and scalability (Fig. 5a). In contrast, RapidNJ fails to scale beyond 200,000 sequences and requires over 23.9-fold more time at that scale. DIPPER’s speed and scalability also facilitates iterative alignment-phylogeny co-estimation, a process that improves tree and alignment quality over successive iterations (Fig. 5b). For example, across three iterations on a one-million-taxon dataset, the tree error decreases from 23.1% to 17.1%, and the alignment error decreases from 5.3% to 1.5% (Fig. 5c), demonstrating that DIPPER can be combined with state-of-the-art progressive aligners, like TWILIGHT, to co-estimate high-quality phylogenies and alignments at scale. It is worth noting that though DIPPER has slightly better accuracy in tree estimation compared to RapidNJ across all iterations, it did not impact the alignment accuracy (Fig. 5b-c), suggesting that MSA inference is resilient to minor errors in tree inference. DIPPER also produces substantially lower tree errors at high speedups compared to recently-proposed “clustering” heuristics for scalable guide tree construction, such as the *K-medoids* clustering method used in FAMSA-2^56^ (Supplementary Fig. 2). Despite the high speedup enabled through DIPPER, given the massive acceleration for MSA estimation recently enabled with TWILIGHT, guide tree construction remains the dominant step in the overall process, constituting up to 82% of the total runtime.

**Figure 5:**
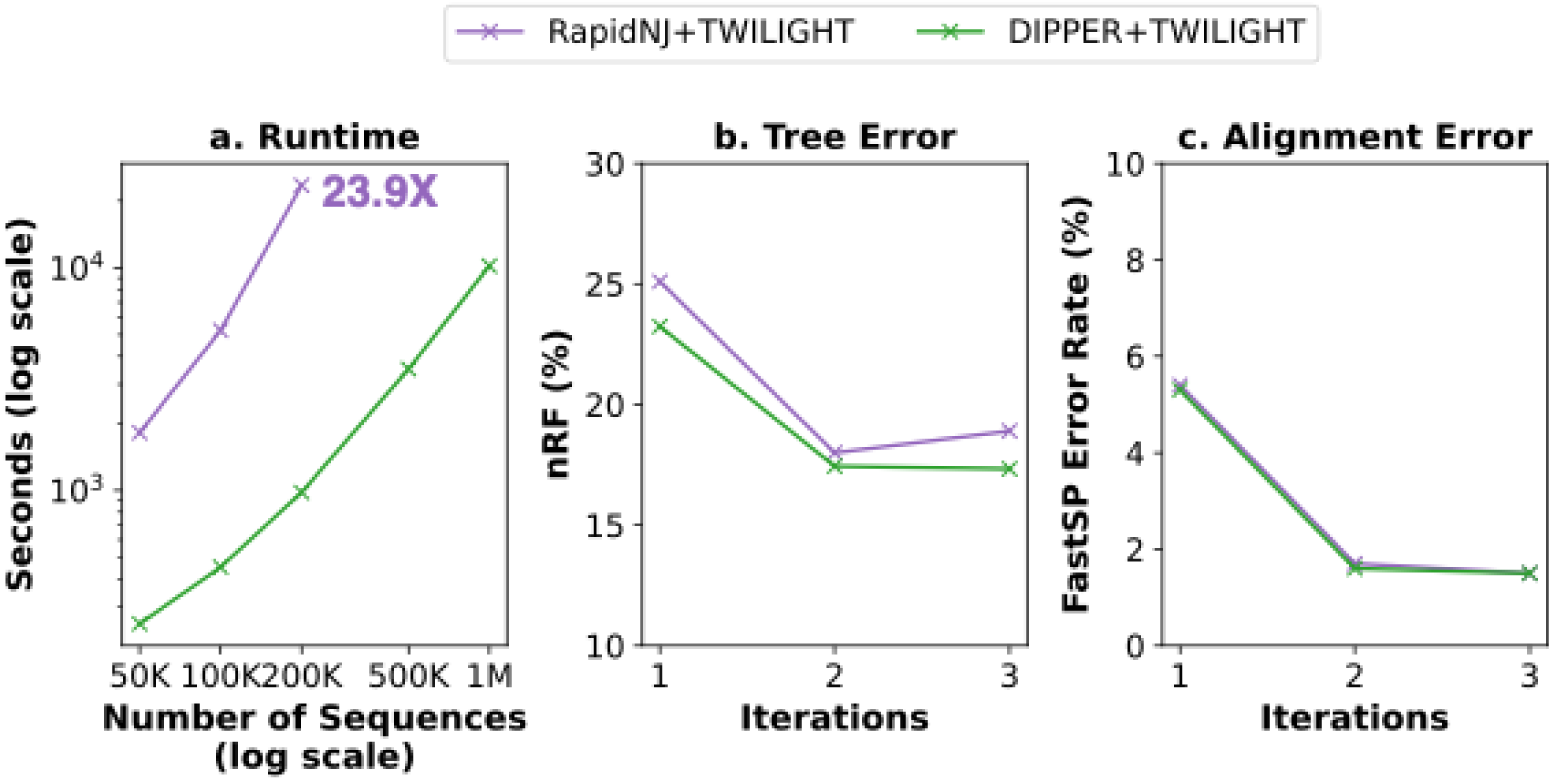
Utility of DIPPER in ultralarge alignment construction. **(a)** Comparison of runtime to generate MSA from raw sequences, generated by AliSim, using TWILIGHT through guide tree construction using DIPPER and RapidNJ. **(b)** Tree error rate and **(c)** Alignment error rate over multiple iterations of tree-alignment co-estimation with DIPPER+TWILIGHT and RapidNJ+TWILIGHT for 200,000 taxa.

### DIPPER enables scalable phylogenetic placement with high accuracy

The ability to place query sequences onto a pre-existing phylogenetic tree, commonly referred to as phylogenetic placement, has emerged as a powerful alternative to *de novo* tree reconstruction, particularly in large-scale analyses where datasets are growing rapidly and computational efficiency is critical. This approach has found widespread use in diverse applications, including microbiome profiling^57^, genome skimming^58^, and pathogen surveillance^59^, where rapid contextualization of new sequences is essential. A principal advantage of phylogenetic placement lies in its scalability: by leveraging a fixed reference topology, placement can be performed orders of magnitude faster than *de novo* inference^60^.

DIPPER provides functionality that enables efficient addition of query sequences onto a provided backbone tree. To evaluate the accuracy and scalability of this functionality, we used simulated datasets generated with RNASim, ranging in size from 20,000 to 1,000,000 tips. For each dataset, we randomly selected 10,000 aligned sequences to construct a backbone tree and placed the remaining sequences onto it. We used DIPPER’s phylogenetic placement functionality to construct the backbone tree as well as to place the query sequences. We compared DIPPER’s performance with APPLES-2^40^ (Fig. 6), a state-of-the-art distance-based phylogenetic placement tool. For APPLES-2, backbone trees were constructed using DecentTree, followed by re-estimation of branch lengths with FastTree^61,62^.

**Figure 6:**
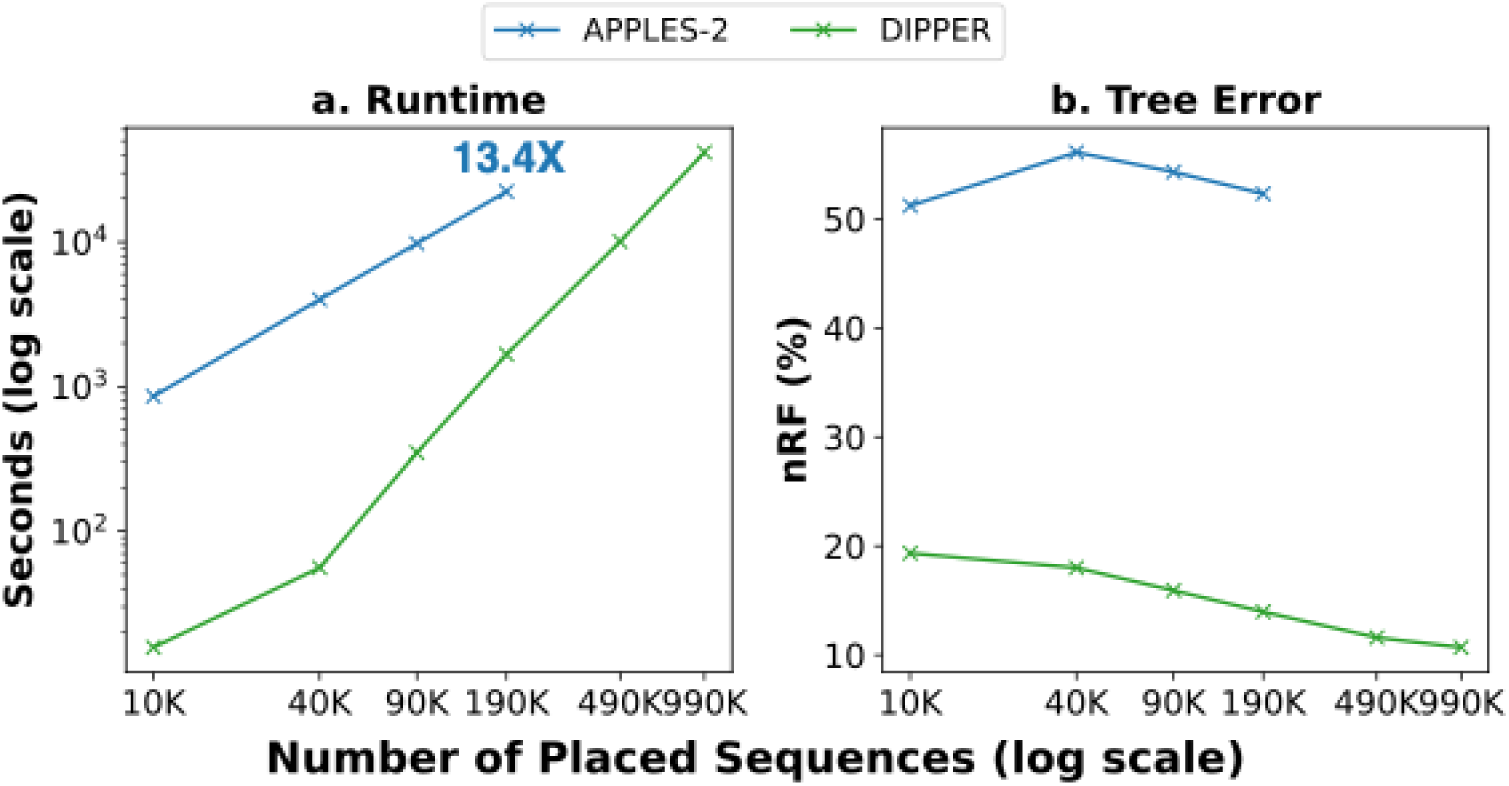
Performance of DIPPER’s phylogenetic placement. Comparison of **(a)** runtime and **(b)** tree error rate of DIPPER with APPLES-2 through adding query sequences to a backbone tree containing 10,000 taxa.

DIPPER demonstrates superior performance in speed, scalability, and accuracy compared to APPLES-2 (Fig. 6). While APPLES-2 failed to place 490,000 query sequences onto the backbone tree within a 24-hour time limit, DIPPER successfully placed 990,000 sequences in just 11 hours and 45 minutes. For a dataset of 190,000 query sequences, DIPPER achieved a 13.4-fold speedup over APPLES-2. In addition to its computational advantages, DIPPER consistently outperformed APPLES-2 in placement accuracy, particularly achieving 41% lower error rate than APPLES-2 after placing 190,000 samples on the backbone tree. Across all tested dataset sizes, DIPPER maintained a lower error rate, which further improved as the number of placed queries increased. Specifically, the error rate decreased from 19.3% to 10.7% as the number of query sequences grew from 10,000 to 190,000.

These results highlight DIPPER’s ability to handle large-scale datasets with both speed and precision, making it a practical choice for high-throughput phylogenetic placement tasks. Its performance enables routine placement of hundreds of thousands of sequences within a single day, unlocking new possibilities for real-time pathogen surveillance, metagenomic analysis, and large-scale evolutionary studies.

### DIPPER minimizes underestimations in the tree path lengths

Distance-based methods such as NJ, UPGMA, and ME, are known to be statistically consistent when provided with an additive distance matrix^7^. However, real-world biological datasets rarely satisfy this requirement. As a result, most phylogenetic tools attempt to fit non-additive distance matrices by optimizing branch lengths to minimize deviations from the input distance estimates, often using least-squares criteria^38,40^. Consequently, path lengths between taxa in the inferred trees often deviate in both directions from the input distances, with roughly half being over- or underestimates (Table 1).

**Table 1:** Percentage of underestimations. (i.e., when a pair of taxa has path length in the inferred tree shorter than the corresponding distance estimate) incurred by baseline tools and DIPPER in its default mode (*K*=10) and strict mode (*K*=-1).

| Tools | CCphylo | RapidNJ | QuickTree | DecentTree | FastME | DIPPER<br>($K=10$, default) | DIPPER<br>($K=-1$, strict mode) |
| --- | --- | --- | --- | --- | --- | --- | --- |
| Underestimations(%) | 52.50% | 50.45% | 50.60% | 50.60% | 50.45% | 2.91% | 0% |

Prior studies have shown that both pairwise alignment methods and commonly used evolutionary models (e.g., JC69^48^, HKY^63^) tend to *underestimate* evolutionary distances, particularly at high divergence levels^64,65^. This skew can also hamper phylogenetic inference^62^. In contrast to prior approaches, DIPPER adopts a novel branch length estimation strategy during placement that explicitly avoids underestimating path lengths relative to the distances to each query’s *K* nearest neighbors. In default mode, DIPPER reduces underestimations to under 3%, compared to a little over 50% in other tools (Table 1). Additionally, DIPPER offers a strict mode (setting *K*=–1) that entirely eliminates underestimation by considering all existing taxa in the tree, not just the local neighborhood. While this increases the runtime from *O(N.*log(*N))* to *O(N*^2^), it can be valuable when distance estimates are likely to be lower bounds on the true evolutionary divergence.

We believe that this unique property of DIPPER to minimize path-length underestimation is not only helpful in practical scenarios, but also results in improved accuracy over previous formulations.vio

### DIPPER’s divide-and-conquer strategy enables the construction of phylogeny with 10 million taxa

DIPPER’s *K-closest placement algorithm,* coupled with efficient GPU implementation, enables it to scale well for datasets with up to one million taxa. Beyond this scale, DIPPER employs a divide-and-conquer strategy that decomposes the problem and applies the placement algorithm to substantially smaller subtrees, using a constant backbone of size. To estimate the impact of the divide-and-conquer strategy on inference accuracy, we set the backbone size to *N/100*, where *N* is the total number of taxa that we varied between 200K to 10M. On datasets up to 2M taxa, up to which the placement strategy scales within practical limits, DIPPER’s divide-and-conquer strategy achieves an 8.1-fold speed-up over its placement strategy (Fig. 7). This implies that compared to its placement strategy, DIPPER’s divide-and-conquer strategy results in an increase in error rate (from 20.8% to 28.2%), which may be acceptable for applications that are tolerant to minor tree inference inaccuracies, such as serving as input guide tree for ultralarge MSAs using tools like TWILIGHT or as an initial tree for scalable tree optimization methods like matOptimize^66^. The divide-and-conquer strategy also allows for unprecedented scale, as DIPPER could reconstruct a phylogeny of 10 million taxa–to our knowledge, the largest for a distance-based phylogenetic tool–in just 6 hours and 28 minutes (Fig. 7). The improved scalability, speed, and high overall accuracy of DIPPER could unlock new applications at previously unattainable scales.

**Figure 7:**
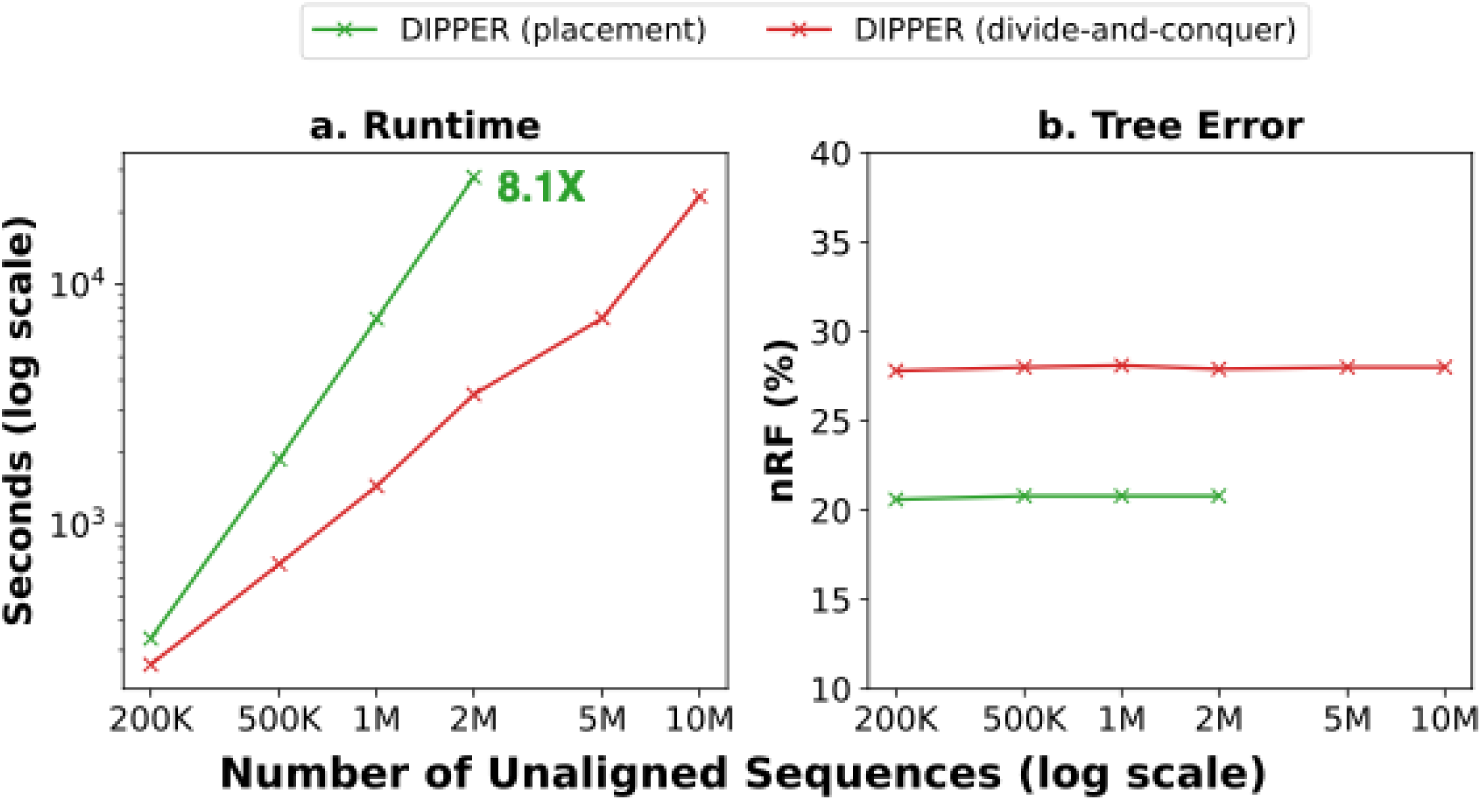
Demonstration of DIPPER’s divide-and-conquer scalability. (a) Runtime and (b) tree error rate of DIPPER using *placement algorithm* (*K*=10) and divide-and-conquer strategy.

## Conclusion and Future Work

In this work, we introduced DIPPER, a distance-based phylogenetic placement tool, designed to address the scalability limitations of existing methods. Leveraging GPU acceleration, DIPPER enables rapid phylogeny reconstruction from both aligned and unaligned sequences, as well as directly from distance matrices. Our experimental results demonstrate that DIPPER achieves substantial speedups over state-of-the-art tools, while maintaining comparable or even superior accuracy.

DIPPER’s efficiency stems from a combination of divide-and-conquer strategies, a novel placement algorithm, memory-efficient design, and parallel computing techniques. These innovations allow it to handle ultralarge datasets with ease, making it a practical and accessible solution for researchers working with ever-growing genomic datasets. Notably, DIPPER is the first tool capable of reconstructing a phylogeny comprising 10 million taxa within hours, marking a significant milestone in computational phylogenetics.

As the volume of genomic data continues to expand exponentially, DIPPER is well-positioned to become a transformative tool in large-scale phylogenetic analyses. Future enhancements may include support for protein sequences and integration of tree refinement techniques, such as Subtree Pruning and Regrafting (SPR)^67^ and Nearest Neighbor Interchange (NNI)^68^, to further improve tree quality. Additionally, support for multi-GPU and distributed high-performance computing (HPC) architectures could further accelerate DIPPER’s performance, and adapting its core innovations for maximum likelihood-based phylogenetics also presents an exciting direction for future research.

## Acknowledgements

We thank Anoushka Saraswat and Arnav Saxena for their initial prototyping efforts, which were helpful in the later stages of DIPPER’s code development. We thank all members of the Turakhia Lab for helpful feedback.

