## Supplementary Figure for "Ultrafast and Ultralarge Distance-Based Phylogenetics Using DIPPER"

### Supplementary Material

#### Appendix A.1: Runtime and Space Complexity Analysis of the Divide-and-Conquer Strategy

**Notation:**
Total taxa: *N*
Maximum backbone and cluster sizes (user-specified constant): *B*
*K* (user-specified constant): used in the *K-closest* heuristic

**Runtime complexity:**

1) Backbone tree: Building a starting backbone tree of size *B* using the placement strategy requires $O(B^2)$ time.

2) Clustering sequences (recursive) and partial tree construction: In the divide stage, DIPPER assigns each of the $N - B$ remaining taxa to one of $2B - 3$ clusters since the backbone tree (which is unrooted) has $2B - 3$ edges. Therefore, the initial clustering has a runtime complexity of $O(NB)$.

At the end of the first clustering step, each cluster has been assigned an expected number of taxa equal to $\frac{N-B}{2B-3}$, or roughly $\frac{N}{2B}$ (since *N*>>>*B*>>*3*). If this value exceeds *B*, each cluster would be recursively subdivided till each cluster has a maximum of *B* new taxa assigned.

After *j* recursive levels, the average cluster size is $\frac{N}{(2B)^j}$ since each level downsizes the cluster size by an expected factor of *2B*. Therefore, the expected number of recursion levels, *J*, before the clusters are smaller than *B* is such that:

$$\frac{N}{(2B)^J} > B$$

$$J > log_{2B}\,(2N) - 1$$

In other words, the expected number of recursion levels scales as $O(log_B\,(N))$. In DIPPER, we limit the number of recursion levels to 2, since we expect the total number of taxa *N* to be within tens to hundreds of millions ($10^7$–$10^8$), and the backbone (and max. cluster) size *B* can be reasonably set to values in the range $10^4$–$10^5$ within practical limits.

In summary, the total runtime of the recursive clustering (divide stage) is $O(N.B.log_B\,(N))$, since there are $O(log_B\,(N))$ recursive levels leading to $O(\frac{N}{B})$ total clusters, each requiring $O(B^2)$ time to build.

3) <u>Merge (conquer) step:</u> In the final stage, the $O(\frac{N}{B})$ total clusters are merged to a final phylogeny of *N* taxa. This is a trivial step with a time complexity of $O(\frac{N}{B})$.

As a result, the total time complexity of the divide-and-conquer strategy is $O(B^2 + N.B.log_B\,(N)\; + \frac{N}{B})$. Since $B$ is a user-specified constant, the expected runtime scales as $O(N.\,log\,(N))$.

The worst-case complexity is still $O(N^2)$ (same as the placement strategy), as there are $O(\frac{N}{B})$ worst-case recursion levels in the divide step.

**Space complexity:**

<u>CPU:</u> The CPU maintains the sketches of all *N* taxa, and also performs the final merge step. Therefore, the space complexity is $O(N)$.

<u>GPU:</u> On GPUs, DIPPER only performs the inference of the partial trees in each cluster (Fig. 1B). Since each cluster has a maximum of *B* taxa, and since this step is performed using the placement strategy which computes an $O(B)$ distance vector on-the-fly for each new placement, its space complexity scales as $O(B)$. In other words, the GPU memory usage is constant (user-specified), independent of the number of taxa.

#### Appendix A.2: Software Versions and Commands Used

| Tool | Version | Command |
|---|---|---|
| **DIPPER** | **v0.1.0** | #Placement<br>dipper -i r -m 1 -I <unaligned sequences> -O <output><br><br>dipper -i m -m 1 -I <aligned sequences> -O <output><br><br>dipper -i d -m 1 -I <distance matrix> -O <output><br><br>#Divide-and-Conquer<br>dipper -i r -m 3 -I <unaligned sequences> -O <output><br><br>dipper -i m -m 3 -I <aligned sequences> -O <output><br><br>dipper -i d -m 3 -I <distance matrix> -O <output> |
| **QuickTree** | **2.5** | quicktree -in a <aligned sequences> > <output><br><br>quicktree -in m <distance matrix> > <output> |
| **RapidNJ** | **2.3.3** | rapidnj <aligned sequences> -i sth -c 32 -a jc > |

| | | |
|---|---|---|
| | | <output><br><br>rapidnj <distance matrix> -i pd -c 32 > <output> |
| **CCPhylo** | **0.8.4** | ccphylo dist -i <aligned sequences> -o ccphylo_dist.phy -t 32 -C 1 && ccphylo tree -i ccphylo_dist.phy -o <output> -t 32<br><br>ccphylo tree -i <distance matrix> -o <output> -t 32 |
| **DecentTree** | **1.0.0** | decenttree -fasta <aligned sequences> -nt 32 -out <output> -t NJ-V -no-banner<br><br>decenttree -in <distance matrix> -nt 32 -out <output> -t NJ-V -no-banner |
| **FastME** | **2.1.6.1** | fastme -i <aligned sequences> -o <output> -m N -T 32 -d J -w n<br><br>fastme -i <distance matrix> -o <output> -m N -T 32 |
| **FastPhylo** | **1.0.0** | fastdist <aligned sequences> -D JC \| fnj -O newick > <output> |
| **APPLES-2** | **2.0.11** | run_apples.py -s <backbone aligned sequences> -x <aligned sequences> -t <backbone tree> -T 32 -D -o apples.jplace --exclude 2> <output> |
| **FAMSA-2** | **2.5.0** | famsa -keep-duplicates -v -medoidtree -t 32 -gt_export <unaligned sequences> <output> |
| **MASH** | **2.2** | mash triangle -p 32 -k 1000 -s 15 <unaligned sequences > <output> |
| **FastTree** | **2.1.11** | fasttree -nome -mllen -intree <backbone tree> -nt <backbone aligned sequences> > <output> |
| **TWILIGHT** | **0.1.3** | twilight -C 32 –G 1 -i <unaligned sequences> -t <input tree> -o <output> --psgop y |

#### Supplementary Figures

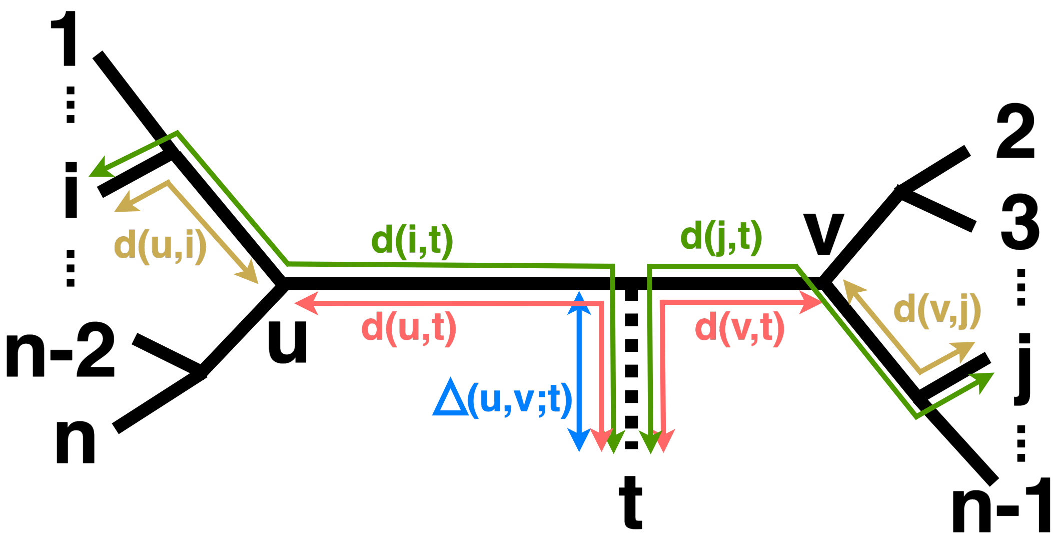


**Supplementary Figure 1:** Illustration of the various notations in DIPPER's placement algorithm. A query taxon *t* is placed on a candidate branch *(u, v)* of the backbone tree with *n* taxa, resulting in additional branch length $\Delta(u, v; t)$.

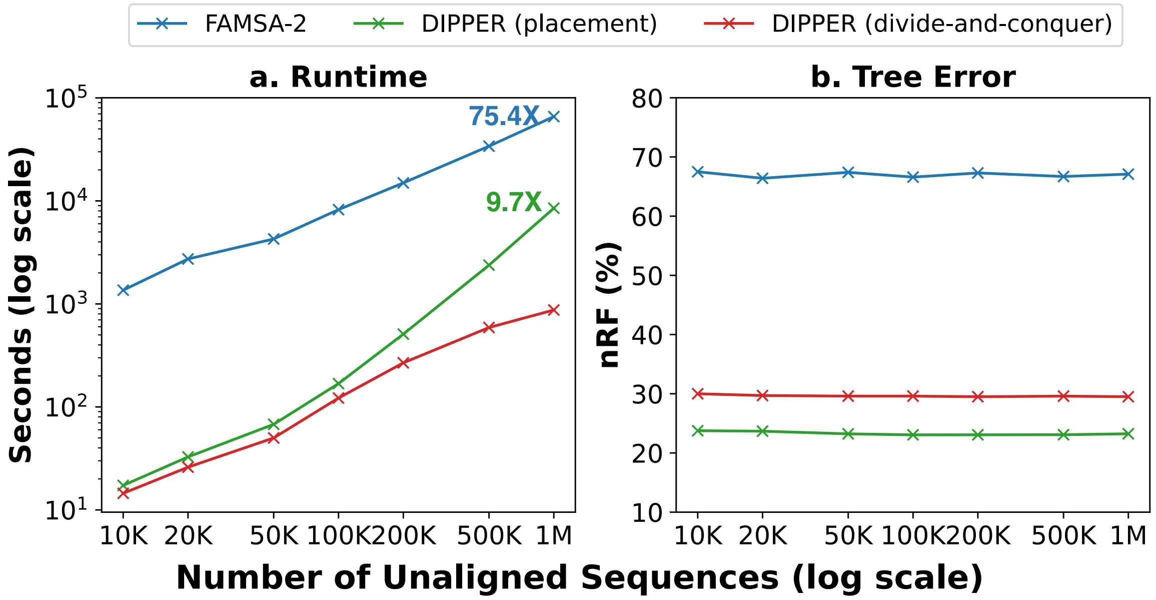


**Supplementary Figure 2: Runtime and accuracy with unaligned sequence count. (a)** Runtime and **(b)** Accuracy of DIPPER (divide-and-conquer) compared with DIPER (placement) and FAMSA-2 to construct phylogenies from unaligned sequences generated using Alisim. We provide the ratio of the runtimes relative to DIPPER (divide-and-conquer).
